# *Verticillium dahliae* strains that infect the same host plant display highly divergent effector catalogs

**DOI:** 10.1101/528729

**Authors:** Hesham A.Y. Gibriel, Jinling Li, Longfu Zhu, Michael F. Seidl, Bart P.H.J. Thomma

## Abstract

During host colonization, plant pathogens secrete molecules that enable host colonization, also known as effector proteins. Here, we show that strains of the fungal plant pathogen *Verticillium dahliae* that are able to infect the same host plant harbour highly divergent LS effector repertoires. Our study outlines the variability within LS effector gene repertoires of *V. dahliae* strains, which may allow the various strains to be competitive in the co-evolution with their hosts.

**Summary:** Effectors are proteins secreted by pathogens to support colonization of host plants, often by deregulating host immunity. Effector genes are often localized within dynamic lineage-specific (LS) genomic regions, allowing rapid evolution of effector catalogues. Such localization permits pathogens to be competitive in the co-evolutionary arms races with their hosts. For a broad host-range pathogen such as *Verticillium dahliae* it is unclear to what extent single members of their total effector repertoires contribute to disease development on multiple hosts. Here, we determined the core and LS effector repertoires of a collection of *V. dahliae* strains, as well as the ability of these strains to infect a range of plant species comprising tomato, cotton, *Nicotiana benthamiana*, Arabidopsis, and sunflower to assess whether the presence of particular LS effectors correlates with the ability to infect particular plant species. Surprisingly, we found that *V. dahliae* strains that are able to infect the same host plant harbor highly divergent LS effector repertoires. Furthermore, we observed differential *V. dahliae* core effector gene expression between host plants. Our data suggest that different *V. dahliae* lineages utilise divergent effector catalogs to colonize the same host plant, suggesting considerable redundancy among the activities of effector catalogs between lineages.

## Introduction

Plant pathogens cause devastating diseases on crop plants, threatening food security worldwide (Fisher et al., 2012; Pennisi, 2010). In order to establish their infection, pathogens secrete effector molecules that can modulate host physiology, often by deregulating host immune responses (Cook et al., 2015; Jones and Dangl, 2006). However, in turn, effectors may become recognized by plant immune receptors, leading to the activation of immune responses and attempted arrest of pathogen invasion (Cook et al., 2015; Jones and Dangl, 2006). Thus, pathogens need to continuously evolve their effector catalogues by modifying or purging existing effectors that became recognized, or by acquiring novel effectors to suppress effector-triggered immune responses (Cook et al., 2015; Jones and Dangl, 2006).

Genomes of plant pathogens are often thought to have evolved two distinct compartments; one comprising gene-rich, repeat-poor genomic regions that contain core genes that mediate general physiology, and one comprising gene-poor, plastic, repeat-rich genomic regions that contain effector genes and other pathogenicity-related genes (Dong et al., 2015; Raffaele and Kamoun, 2012). The plastic genomic regions are either embedded within core chromosomes or reside on separate chromosomes that are often referred to as conditionally dispensable chromosomes (CDCs) (Dong et al., 2015; Raffaele and Kamoun, 2012). For instance, effector genes of the tomato-pathogen *Fusarium oxysporum* f. sp. *lycopersici*, known as secreted in xylem (*SIX*) genes, are located on dispensable chromosomes (Schmidt et al., 2013). In contrast, all known effectors of the fungal wheat pathogen *Zymoseptoria tritici* are located on core chromosomes, while no recognizable effector genes reside on dispensable chromosomes (Kema et al., 2018; Marshall et al., 2011; Meile et al., 2018). Core chromosomes also carry effector genes in other fungal plant pathogens, such as the fungal smut pathogen *Ustilago maydis* and the fungal tomato pathogen *Cladosporium fulvum* (Hemetsberger et al., 2015; Stergiopoulos et al., 2010). A genome compartmentalization with physically separated effector-containing regions is often referred to as a “two-speed” genome organization because it is thought that gene-rich, repeat-poor genomic regions evolve slowly, while gene-poor, repeat-rich genomic regions evolve quicker (Croll and McDonald, 2012; Raffaele and Kamoun, 2012). Accordingly, the occurrence of effector genes within plastic genomic regions allows rapid evolution of effector catalogues and permits pathogens to be competitive in the co-evolution with hosts and evade their immune systems (Dong et al., 2015; Raffaele and Kamoun, 2012).

*Verticillium dahliae* is a soil-borne fungal plant pathogen that is able to infect a broad range of plant species, including crops such as tomato, potato, lettuce, and cotton (Fradin and Thomma, 2006; Inderbitzin et al., 2011). The fungus infects plants through their roots and subsequently colonizes the water-conducting xylem vessels, leading to vascular wilt disease (Fradin and Thomma, 2006). Comparative genomics between closely related *V. dahliae* strains revealed that they carry highly dynamic, repeat rich, lineage-specific (LS) regions that are only present in a subset of *V. dahliae* strains, and that account for up to 4 Mb of the ∼35 Mb genome (de Jonge et al., 2013; Faino et al., 2016). These LS regions are enriched for *in planta-*induced effector genes that contribute to fungal virulence (de Jonge et al., 2013). However, effector genes are not only found in LS regions, as also the core genome harbors effector genes, such as those encoding a family of necrosis and ethylene-inducing-like proteins (NLPs) some of which were found to induce cell death in dicotyledonous plants (de Jonge et al., 2011; Santhanam et al., 2013). Similarly, also a family of lysin motif (LysM) effectors is encoded in the core genome, various homologs of which have been reported to enhance virulence by suppression of chitin-triggered immunity in other fungal pathogens (de Jonge et al., 2010; Kombrink et al., 2017; Marshall et al., 2011; Mentlak et al., 2012; Takahara et al., 2016). However, only a single LysM effector that is encoded in an LS region of *V. dahliae* strain VdLs17, and is thus not identified in the genomes of other *V. dahliae* strains, was found to contribute to virulence by suppression of chitin-triggered immunity, whereas no role in virulence could be attributed to any of the core LysM effectors (Kombrink et al., 2017).

Whereas *V. dahliae* is characterized by its generally broad host range, differential pathogenicity among hosts occurs for individual strains (Bhat and Subbarao, 1999). In this study, we analysed the genomes of a collection of *V. dahliae* strains and assessed their core and LS effector catalogues in relation to their host ranges. To this end, we selected a set of strains that are well-adapted to cause disease on tomato (*Solanum lycopersicum*), cotton (*Gossypium hirsutum*), Australian tobacco (*Nicotiana benthamiana*), Arabidopsis (*Arabidopsis thaliana*), and sunflower (*Helianthus annuus*) and determined their core and LS effector catalogues as well as their *in planta* expression profiles.

## Results

### *Verticillium dahliae* pathogenicity on a panel of potential host plants

To evaluate the pathogenicity of a collection of *V. dahliae* strains on a panel of potential host plant species, we conducted inoculation experiments with 21 strains on the Solanaceae crop plant tomato and model plant *N. benthamiana*, the Malvaceae crop plant cotton, the Asteraceae crop plant sunflower, and the Brassicaceae model plant Arabidopsis. Despite the fact that *V. dahliae* is often considered a broad host range pathogen, there is no individual *V. dahliae* strain in this collection that is able to cause disease on all tested plant species (Table 1). All isolates are pathogenic on Arabidopsis (Figure 1) and on *N. benthamiana* (Figure 2), albeit that the severity of disease symptoms induced by different strains vary considerably (Figure 1 & Figure 2). Most strains were also found to cause disease on cotton, with the exception of strains 2009-605 and V152 that are non-pathogenic on this host species (Figure 3). Interestingly, *V. dahliae* strains CQ2, 463 and ST100 cause severe defoliation, while the other pathogenic strains induce mild to moderate disease symptoms that include wilting, stunting and chlorosis in the absence of defoliation (Figure 3). Fewer *V. dahliae* strains are able to cause disease on tomato (Figure 4). Besides several strains that cause defoliation on cotton, the tomato non-pathogenic strains also include several non-defoliators on cotton like Vd39 and 85S (Figure 3 & Figure 4). Interestingly, whereas strain V152 that is non-pathogenic on cotton also fails to cause disease on tomato, strain 2009-605 that is non-pathogenic on cotton is able to cause wilt disease on tomato (Figure 4). Strikingly, except for *V. dahliae* strain 85S that induces clear wilt disease symptoms on sunflower plants, including stunting, chlorosis and necrosis, all other strains fail to cause disease on sunflower (Figure 5). Thus, differential pathogenicity occurs within the collection of *V. dahliae* strains tested here.

**Table 1.** Inoculation experiments with a collection of *V. dahliae* strains on a collection of potential host plants

| Strain | Arabidopsis | *N. benthamiana* | Cotton | Tomato | Sunflower |
| --- | --- | --- | --- | --- | --- |
| ST100 | + | + | + | - | - |
| 463 | + | + | + | - | - |
| CQ2 | + | + | + | - | - |
| Vd39 | + | + | + | - | - |
| 85S | + | + | + | - | + |
| JKG8 | + | + | + | + | - |
| DVD-S94 | + | + | + | + | - |
| DVD3 | + | + | + | + | - |
| DVD31 | + | + | + | + | - |
| DVD161 | + | + | + | + | - |
| DVD-S29 | + | + | + | + | - |
| ST14.01 | + | + | + | + | - |
| CBS38166 | + | + | + | + | - |
| DVD-S26 | + | + | + | + | - |
| Vandijk | + | + | + | - | - |
| ST16.01 | + | + | + | - | - |
| 2009-605 | + | + | - | + | - |
| V152 | + | + | - | - | - |
| V52 | + | + | + | + | - |
| VdLs17 | + | + | + | + | - |
| JR2 | + | + | + | + | - |

'+'= pathogenic; '-'= non-pathogenic. Disease symptoms were scored up to 21 days post inoculation (dpi) (tomato, *N. benthamiana*, Arabidopsis), 28 dpi (cotton) or 45 dpi (sunflower). The inoculation experiments were executed twice with similar results.

**Figure 1.**
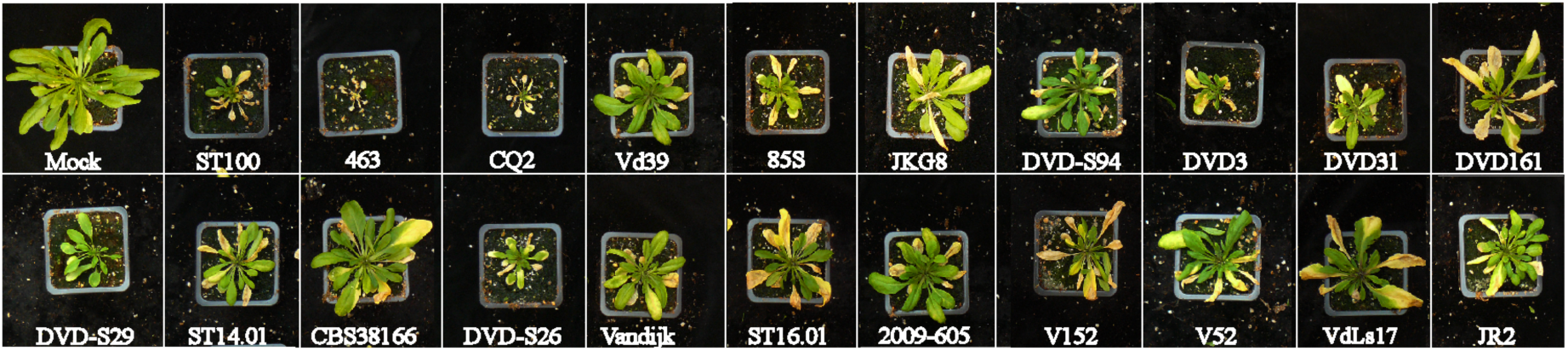
Phenotypes of *Arabidopsis thaliana* inoculated with *V. dahliae* strains. Typical appearance of *A. thaliana* (Col-0) plants upon mock-inoculation or inoculation with a collection of *V*. *dahliae* strains. All inoculated *V. dahliae* strains are pathogenic on *A. thaliana*. Note that disease symptoms range from stunting, wilting to severe tissue necrosis. Pictures show representative plants at 21 days after inoculation taken from one of two independent inoculation experiments.

**Figure 2.**
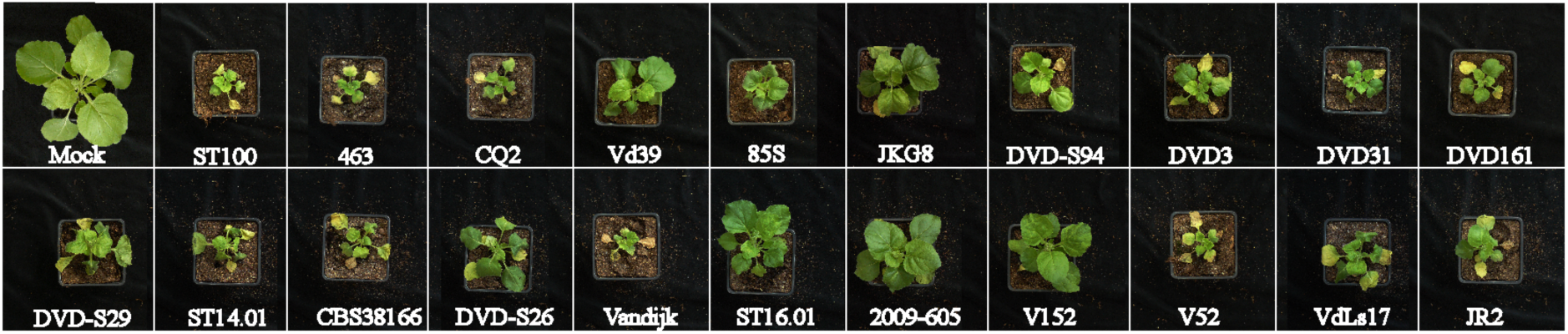
Phenotypes of *Nicotiana benthamiana* inoculated with *V. dahliae* strains. Typical appearance of *N. benthamiana* plants upon mock-inoculation or inoculation with a collection of *V. dahliae* strains. All inoculated *V. dahliae* strains are pathogenic on *N. benthamiana*. Note that disease symptoms include stunting, wilting and severe tissue necrosis. Pictures show representative plants at 21 days after inoculation taken from one of two independent inoculation experiments.

**Figure 3.**
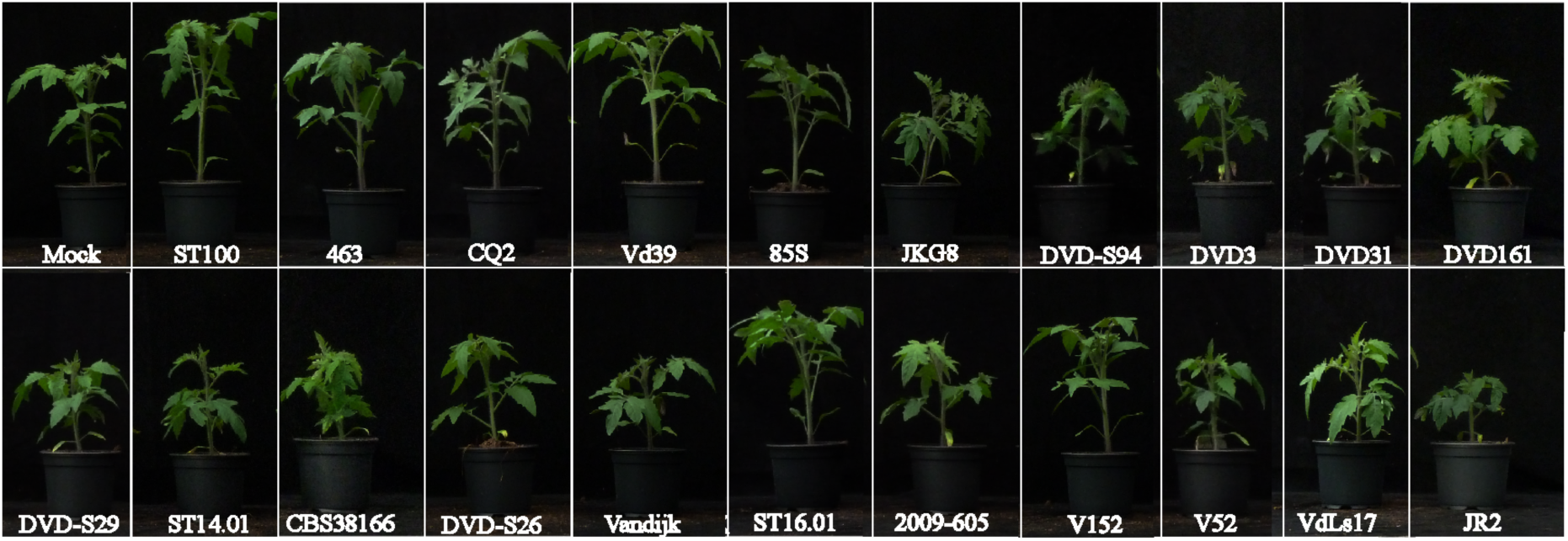
Phenotypes of cotton plants inoculated with *V. dahliae* strains. Typical appearance of cotton (cv. Simian 3) plants upon mock-inoculation or inoculation with a collection of *V. dahliae* strains. *V. dahliae* strains display differential pathogenticity on cotton plants. Note that several *V. dahliae* strains cause defoliation symptoms, while others induce wilting, stunting but not defoliation. Pictures show representative plants at 28 days after inoculation taken from one of two independent inoculation experiments.

**Figure 4.**
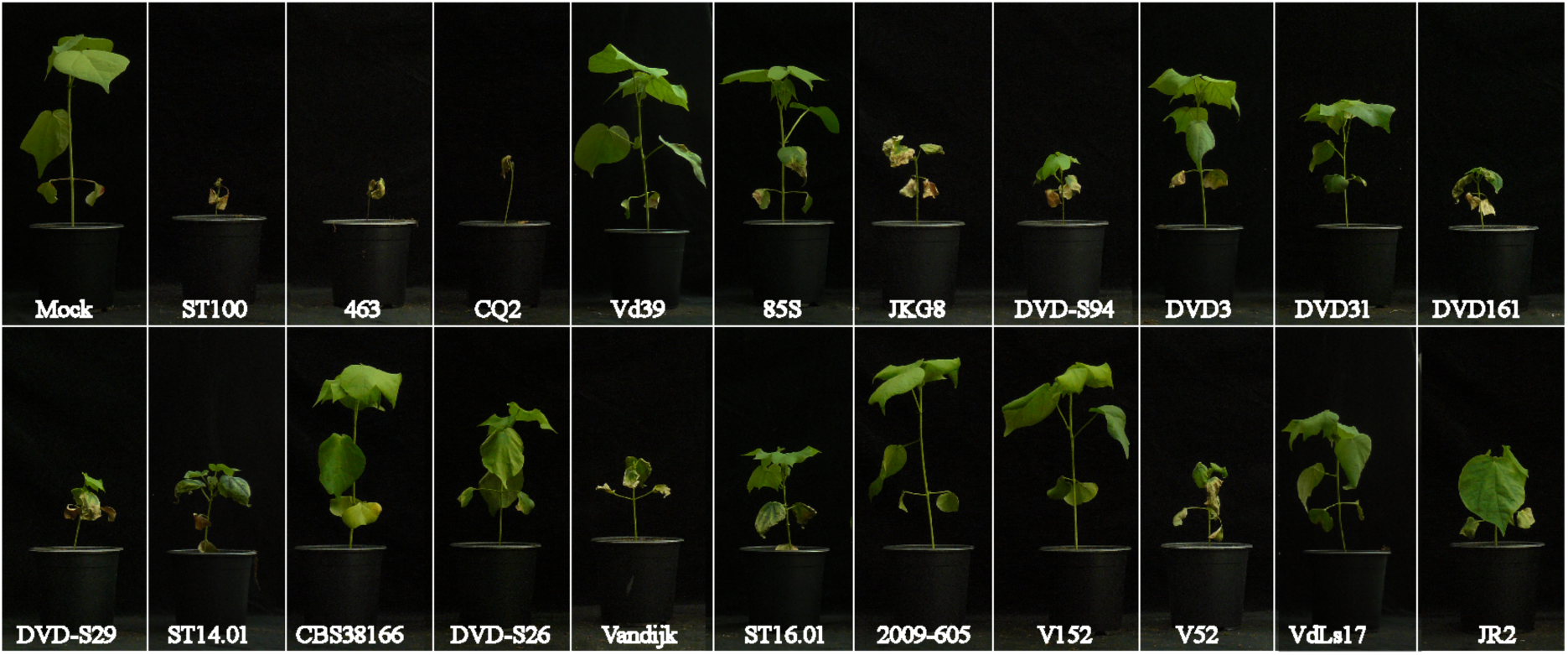
Phenotypes of tomato plants inoculated with *V. dahliae* strains. Typical appearance of tomato (cv. Moneymaker) plants upon mock-inoculation or inoculation with a collection of *V. dahliae* strains. *V. dahliae* strains display differential pathogenticity on tomato plants. Note that pathogenic strains induce clear stunting and significant reduction in canopy area development on inoculated plants. Pictures show representative plants at 21 days after inoculation taken from one of two independent inoculation experiments.

**Figure 5.**
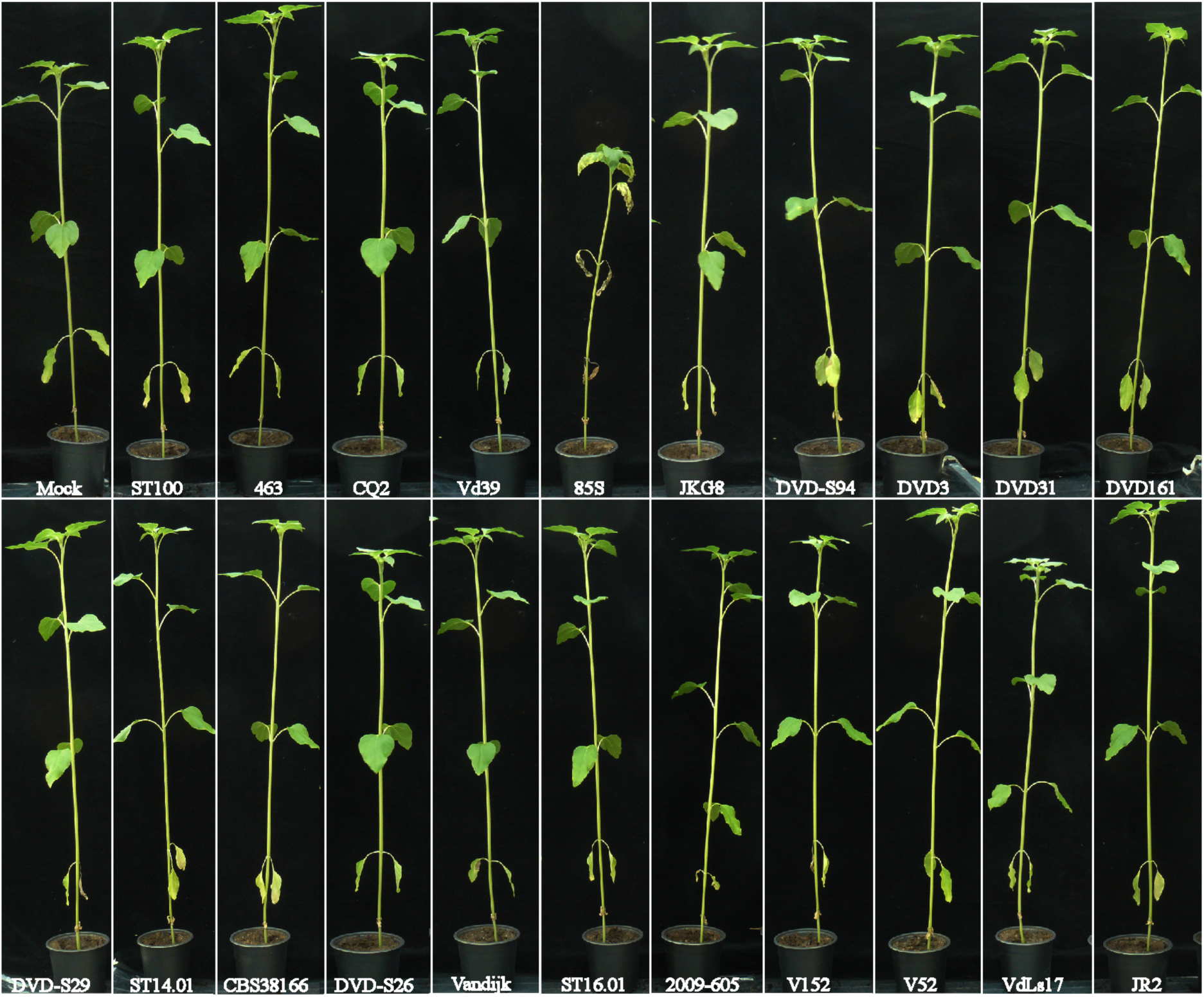
Phenotypes of sunflower plants inoculated with *V. dahliae* strains. Typical appearance of sunflower (cv. Tutti) plants upon mock-inoculation or inoculation with a collection of *V. dahliae* strains. *V. dahliae* strains display differential pathogenticity on sunflower plants Note that besides the stunting, the plant inoculated with pathogenic strain 85S also displays chlorosis and wilting symptoms. Pictures show representative plants at 45 days after inoculation taken from one of two independent inoculation experiments.

### Genome assemblies and annotations of a collection of *V. dahliae* strains

The genome sequences of 13 *V. dahliae* strains (Table S1) were obtained from previous studies, nine of which were determined using the Illumina HiSeq 2000 platform (de Jonge et al., 2013; de Jonge et al., 2012), and four were sequenced using long-read PacBio Single-Molecule Real-Time (SMRT) sequencing technology (Faino et al., 2015). In this study, we sequenced the genomes of eight additional *V. dahliae* strains (Table S1) using the Illumina HiSeq 2000 platform, yielding ∼1.1 Gb of paired-end (PE) library-derived reads (500 bp insert size; 100 bp read length) per strain. As the genomes of *V. dahliae* strains that were previously sequenced with Illumina technology showed a reduced N50 size of about 35.55 kb (de Jonge et al., 2012), all the Illumina sequenced genomes were re-assembled in this study. The short reads of the 17 *V. dahliae* strains that were sequenced with Illumina technology were assembled into ∼34 Mb, with the largest assembly of 35.90 Mb for *V. dahliae* strain Vd39, and the smallest assembly of 33.14 Mb for *V. dahliae* strain DVD-S29 (Table S2). All assemblies comprised between 1,000 and 4,188 scaffolds with an N50 of ∼50 kb, except for *V. dahliae* strains 463 and V52 that were assembled in 4,188 and 3,419 scaffolds with an N50 of 17.74 kb and 21.80 kb, respectively (Table S2).

To assess the completeness of the assemblies, the Benchmarking Universal Single-Copy Orthologs (BUSCO) software was used, which uses a set of 1,315 core Ascomycota genes as queries (Simão et al., 2015). The BUSCO scores amounted to ∼91% for all the assemblies, except for *V. dahliae* strains 463 and v52 that resulted in 75% and 78.6%, respectively (Table S2). For the PacBio sequenced strains, BUSCO scored 99.30% for 85S and 97.50% for CQ2, while the BUSCO scores for the gapless genome assemblies of JR2 and VdLs17 (Faino et al., 2015) amounted to 99.40% and 98.90%, respectively (Table S2).

Repetitive elements are strong drivers of genome evolution in plant pathogens (Seidl and Thomma 2017). Thus, the amounts and types of repetitive elements in the genomes of *V. dahliae* strains were predicted by combining *de novo* and known repetitive elements with RepeatMasker (Smit et al., 2016). The repeat content within the *V. dahliae* genomes varied between 6.64% (2.26 Mb) for *V. dahliae* strain 2009-605 and 13.43% (4.83 Mb) for *V. dahliae* strain 85S (Table S3). Out of all the annotated repetitive elements, different repeat families were identified, which included long terminal repeats (LTRs) (2 Mb, ∼5.7%), long interspersed nuclear elements (LINEs) (40 kb, ∼0.11%), and short interspersed nuclear elements (SINEs) (2.9 kb, ∼0.01%) (Table S3).

We subsequently inferred gene annotations for the various *V. dahliae* strains. For *V. dahliae* strains JR2 and CQ2, a previously determined gene annotation was used (Faino et al., 2015). The completeness of gene annotation for both strains was assessed using BUSCO that scored only a low score for CQ2 (72%) compared to JR2 (90.80%). Thus, gene annotations were inferred for all 20 *V. dahliae* strains, except for *V. dahliae* strain JR2. The Maker2 pipeline (Holt and Yandell, 2011) was used that combines *de novo*, homology-based, and previous gene annotations for *V. dahliae* strains CQ2 (B. Thomma and J. Li, unpublished) and JR2 (Faino et al., 2015), and protein homologs of 260 predicted fungal proteomes. The number of genes varied from 10,461 for *V*. *dahliae* strain VanDijk to 11,341 for *V. dahliae* v52 (Table 2).

**Table 2.** Summary of genome sizes, core and lineage-specific (LS) regions, number of genes, and number of effector genes for each strain.

| Strain | Genome size (Mb) |  |  |  |  |  |  | Number of genes |  |  |  |  |  | EffectorP |  |  |
| --- | --- | --- | --- | --- | --- | --- | --- | --- | --- | --- | --- | --- | --- | --- | --- | --- |
|  | Total | Repeats | %Repeats | Core | %Core | LS | %LS | Total | Secreted | Core | LS | Core/secreted | LS/secreted | Total | Core | LS |
| 2009-605 | 34.06 | 2.26 | 6.64 | 32.16 | 94.43 | 1.90 | 5.57 | 10893 | 1105 | 9707 | 1186 | 1020 | 85 | 190 | 170 | 20 |
| 463 | 34.03 | 3.56 | 10.46 | 32.66 | 95.96 | 1.38 | 4.04 | 11188 | 1002 | 10671 | 517 | 967 | 35 | 204 | 190 | 14 |
| 85S | 35.93 | 4.83 | 13.43 | 34.11 | 94.93 | 1.82 | 5.07 | 10580 | 1062 | 9706 | 874 | 995 | 67 | 174 | 168 | 6 |
| CBS38166 | 34.03 | 2.83 | 8.32 | 32.59 | 95.76 | 1.44 | 4.24 | 10676 | 1024 | 9823 | 853 | 969 | 55 | 180 | 168 | 12 |
| DVD161 | 33.47 | 2.61 | 7.81 | 32.31 | 96.53 | 1.16 | 3.47 | 10624 | 1035 | 9872 | 752 | 980 | 55 | 195 | 172 | 23 |
| DVD31 | 33.58 | 2.85 | 8.50 | 32.48 | 96.72 | 1.10 | 3.28 | 10642 | 1034 | 9941 | 701 | 981 | 53 | 185 | 164 | 21 |
| DVD3 | 34.42 | 3.47 | 10.10 | 33.04 | 96.00 | 1.37 | 4.00 | 10741 | 1035 | 9934 | 807 | 982 | 53 | 191 | 172 | 19 |
| DVD-S26 | 34.74 | 2.82 | 8.11 | 32.59 | 93.82 | 2.15 | 6.18 | 10977 | 1057 | 9871 | 1106 | 979 | 78 | 196 | 178 | 18 |
| DVD-S29 | 33.14 | 2.55 | 7.69 | 32.07 | 96.77 | 1.07 | 3.23 | 10561 | 1020 | 9866 | 697 | 978 | 42 | 193 | 178 | 15 |
| DVD-S94 | 34.42 | 3.31 | 9.61 | 32.98 | 95.80 | 1.44 | 4.20 | 10684 | 1057 | 9827 | 857 | 990 | 67 | 185 | 165 | 20 |
| JKG8 | 33.85 | 2.75 | 8.13 | 32.56 | 96.21 | 1.28 | 3.79 | 10565 | 1074 | 9647 | 918 | 1006 | 68 | 170 | 156 | 14 |
| ST100 | 34.92 | 3.66 | 10.49 | 33.28 | 95.32 | 1.64 | 4.68 | 10697 | 1041 | 9789 | 908 | 972 | 69 | 189 | 170 | 19 |
| ST14.01 | 34.48 | 3.39 | 9.83 | 32.92 | 95.49 | 1.55 | 4.51 | 10660 | 1039 | 9716 | 944 | 976 | 63 | 175 | 162 | 13 |
| ST16.01 | 34.22 | 2.31 | 6.74 | 31.95 | 93.37 | 2.27 | 6.63 | 11005 | 1108 | 9687 | 1318 | 1017 | 91 | 195 | 174 | 21 |
| V152 | 33.98 | 2.79 | 8.22 | 32.21 | 94.82 | 1.76 | 5.18 | 11137 | 1059 | 10103 | 1034 | 988 | 71 | 195 | 175 | 20 |
| V52 | 33.54 | 2.41 | 7.19 | 31.83 | 94.91 | 1.71 | 5.09 | 11341 | 1030 | 10648 | 693 | 980 | 50 | 207 | 188 | 19 |
| VanDijk | 33.17 | 2.34 | 7.06 | 31.97 | 96.39 | 1.20 | 3.61 | 10461 | 1068 | 9591 | 870 | 999 | 69 | 180 | 163 | 17 |
| Vd39 | 35.90 | 4.45 | 12.39 | 33.69 | 93.84 | 2.21 | 6.16 | 10662 | 1074 | 9579 | 1083 | 988 | 86 | 171 | 159 | 12 |
| CQ2 | 35.82 | 4.15 | 11.60 | 34.03 | 95.00 | 1.79 | 5.00 | 10528 | 1064 | 9628 | 900 | 994 | 70 | 169 | 165 | 4 |
| JR2 | 36.15 | 4.26 | 11.77 | 33.68 | 93.15 | 2.47 | 6.85 | 11432 | 1067 | 10328 | 1104 | 1001 | 66 | 212 | 200 | 12 |
| VdLs17 | 35.97 | 4.20 | 11.68 | 33.52 | 93.19 | 2.45 | 6.81 | 10856 | 1090 | 9688 | 1168 | 1000 | 90 | 179 | 165 | 14 |

### Identification of core and lineage-specific (LS) effector catalogs

Initially, the secretomes for each of the *V. dahliae* strains were predicted, identifying between 1,002 secreted proteins for *V. dahliae* strain 463 and 1,108 proteins for *V. dahliae* strain ST16 (Table 2). Subsequently, the machine-learning algorithm of EffectorP (Sperschneider et al., 2016) was used, which identified between 169 effectors for *V. dahliae* strain CQ2 and 212 effectors for *V. dahliae* strain JR2 (Table 2).

Subsequently, we determined the core genome, here defined as regions that are shared by ≥19 *V. dahliae* strains, and LS regions, here defined as regions that are shared by <19 *V. dahliae* strains, for all *V. dahliae* strains. The core regions of all *V. dahliae* strains comprise 32.79 Mb (93-97%) of the genome, while LS regions comprise between 1.06 and 2.47 Mb (3-7%) (Table 2). On average, the core regions of *V. dahliae* strains harbor 9,886 genes, comprising 988 genes that encode secreted proteins, of which 171 were classified as effectors based on EffectorP (Table 2). The LS regions of *V. dahliae* strains harbor between 517 genes for *V. dahliae* strain 463 and 1,318 genes for *V. dahliae* strain ST16 (Table 2). Of these LS genes, 35 genes encode secreted proteins for *V. dahliae* strain 463 and 91 for *V. dahliae* strain ST16, of which ∼15 genes were classified as effectors for each *V. dahliae* strain (Table 2). We tested these predictions on the previously identified LS effector gene of *V. dahliae* strain JR2, namely *Ave1*, which was successfully identified as a LS effector gene (de Jonge et al., 2012) (Figure S1).

To assess general characteristics of core and LS effector genes, features such as their distance to transposable elements (TEs), gene length, inter-genic length, and expression were determined. For all strains, we observed that LS effector genes are shorter in length than core effector genes (Figure 6A, B; Figure S2). The average inter-genic length of LS effector genes is slightly longer (1476 bp) compared to the average inter-genic length of core effector genes (1232 bp) (Figure 6A, B; Figure S3). Moreover, LS effector genes localize closer to TEs than core effector genes, even though this trend is not significant for *V. dahliae* strain JR2 (Figure 6A, B; Figure S4). Finally, LS effector genes of *V. dahliae* strain JR2 were found to be significantly higher expressed *in planta* on *N. benthamiana* than core effector genes (Figure 7A), although no such difference was found between LS and core effector genes of *V. dahliae* strain CQ2 *in planta* on cotton (Figure 7B).

**Figure 6.**
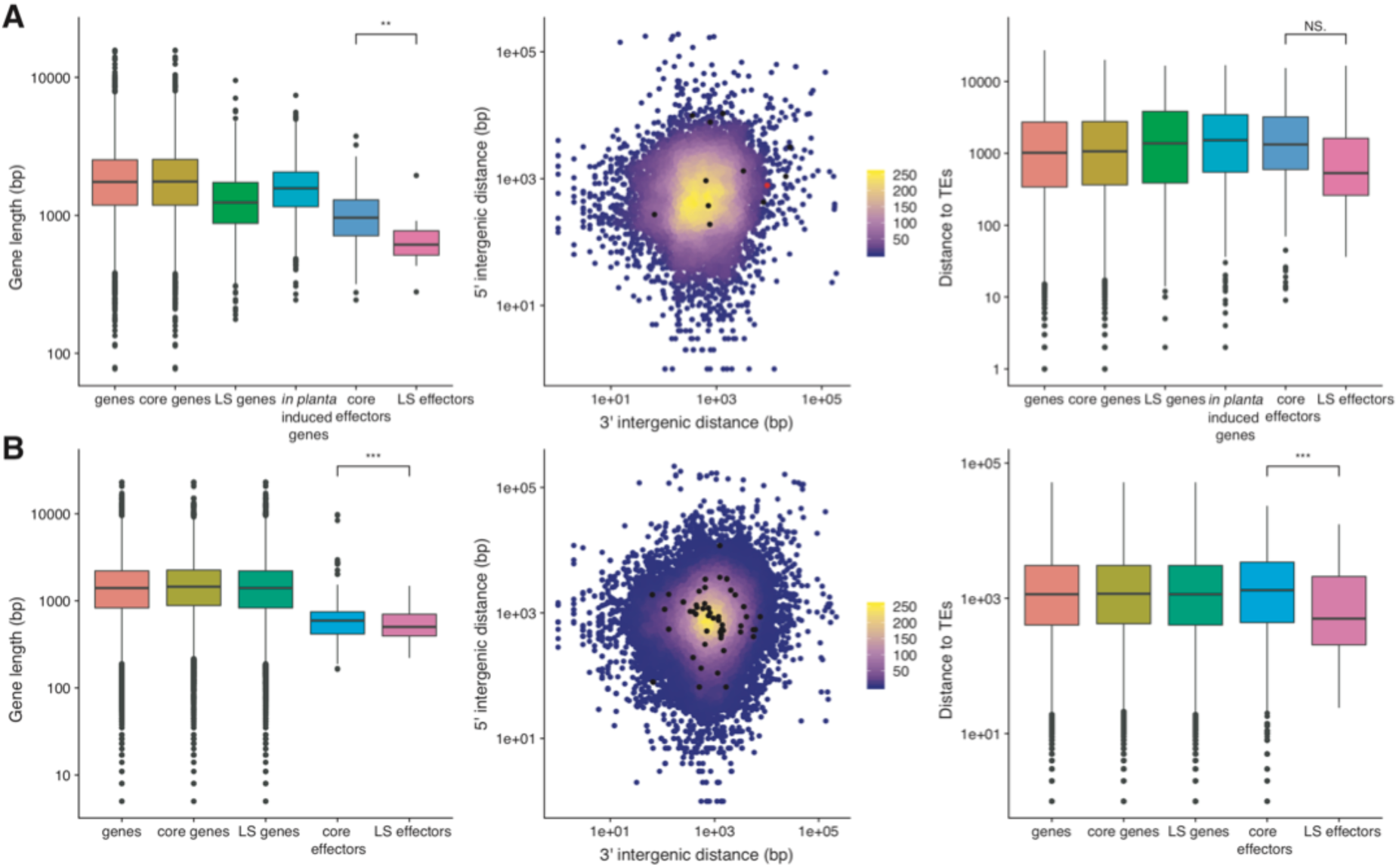
Characteristics of core and LS effector genes. Gene length, two-dimensional density plot of 5′- and 3′-flanking inter-genic regions, where LS effectors are indicated in (black) and Ave1 effector in (red) color, and distance to closest transposable element (TE) for core and LS effector genes of *V. dahliae* strain JR2 **(A)** vs all *V. dahliae* strains **(B)** are displayed.

**Figure 7.**
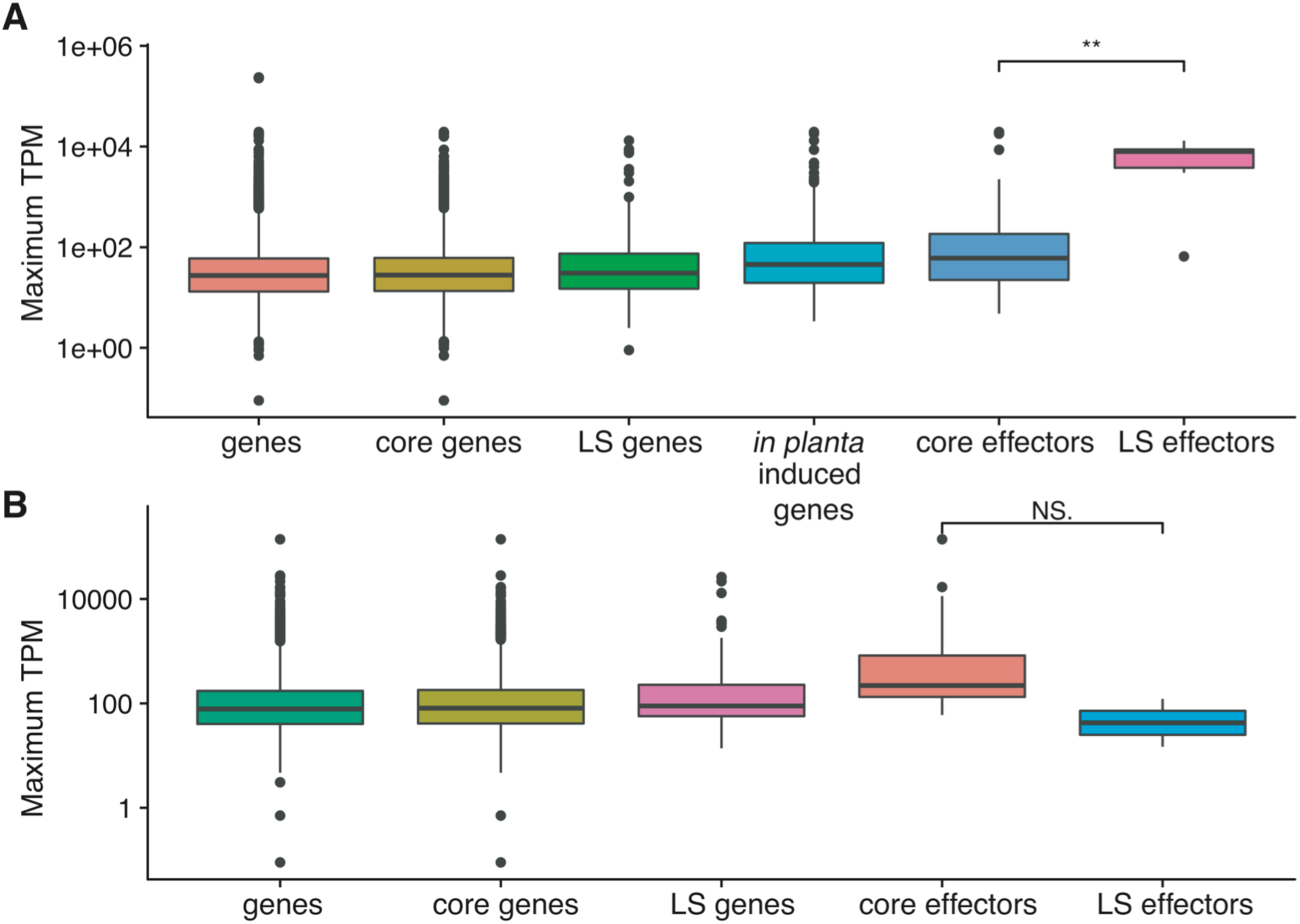
Maximum TPM values of core and LS effector genes over four time points during *N. benthamiana* and cotton infection by *V. dahliae* strains JR2 and V911. **A)** Maximum TPM values per time point (4, 8, 12, and 16) of core and LS effector genes during *N. benthamiana* infection by *V. dahliae* strain JR2. **B)** Maximum TPM values per time point (6, 9, 12, and 15) of core and LS effector genes during cotton infection by *V. dahliae* strain V911.

### *V. dahliae* strains that infect the same host plant harbor divergent effector repertoire

It has previously been shown that *V. dahliae* strains display differential capacity to infect particular host plants (Bhat and Subbarao 1999). We furthermore showed that especially LS effectors contribute to *V. dahliae* pathogenicity on individual plant hosts (de Jonge et al., 2013; de Jonge et al., 2012; Kombrink et al., 2017). Collectively, this suggests that *V. dahliae* strains may harbor an array of specialized effectors that only function on particular host plants. Therefore, we assessed whether the presence of particular LS effectors in the various *V. dahliae* strains correlates with the ability to infect particular hosts. In total, we predicted 333 LS effectors over the various strains (Table 2) that were clustered into 110 families that are either shared by sub-groups of *V. dahliae* strains or are strain-specific (Figure 8). Of these effectors, the previously identified LS effector genes *Ave1* is shared by sub-groups of *V. dahliae* strains (de Jonge et al., 2013), while the *Sun1* effector gene of *V. dahliae* strain 85S is strain specific (Chapter 4) (Figure 8). Notably, in contrast to previous observations, we found that the LS effector genes *XLOC_00170, XLOC_008951*, and *XLOC_009059* were only present in *V. dahliae* strain JR2 and not in additional *V. dahliae* strains (de Jonge et al., 2013). Thus, we searched the 333 LS effector genes against the *V. dahliae* genome assemblies using BLAST (tblastn), which revealed that multiple (candidate) effector genes, including the previous identified ones, were absent from the gene annotation, highlighting the challenge of computational effector gene identification (Gibriel et al., 2016). Intriguingly, we observed highly dissimilar LS effector catalogs among *V. dahlia*e strains that are able to infect the same host plant (Figure 8). Of the 333 LS effectors, not even a single one is shared among strains that infect the same host plant (Figure 8). Even more strikingly, *V. dahliae* strains that infect the same host plant do not cluster based on their LS effector repertoires (Figure 8). Thus, *V. dahliae* strains harbor highly divergent LS effector catalogs, the composition of which does not correlate with the host plant they are able to infect.

**Figure 8.**
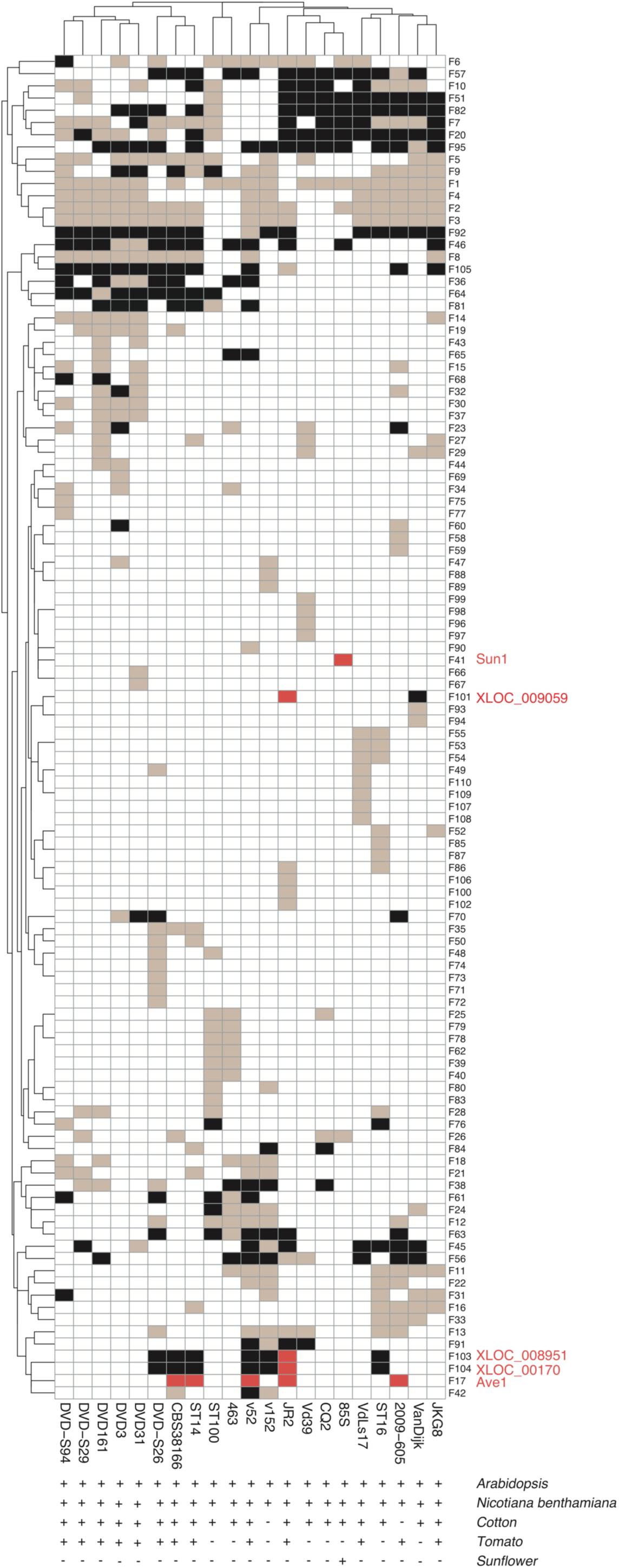
*Verticillium dahliae* strains that infect the same host plant harbor highly divergent LS effector repertoires. LS effectors were clustered into 110 families and their presence (brown) and absence (white) between *V. dahliae* strains is shown. Previously described LS effectors are colored in red (de Jonge et al., 2013; Li et al., unpublished). Black colors indicate the absence of LS effectors from the predicted gene annotation for *V. dahliae* strains and the presence in *V. dahliae* genomes based on BLAST (tblastn).

### Variability in core effector transcription profiles

In many plant pathogens, core effector genes are *in planta* highly*-*induced and play essential roles on a multitude of hosts (Guyon et al., 2014; Hemetsberger et al., 2015; Santhanam et al., 2013; Yin et al., 2017). Thus, we assessed the expression of *V. dahliae* core effector genes during invasion of different plant species. First, we mapped RNA-seq datasets from *N. benthamiana* plants colonized by *V. dahliae* strain JR2 (de Jonge et al., 2012) against the reference genome sequence of *V. dahliae* strain JR2 (Faino et al., 2015), and *G. hirsutum* (cotton) plants colonized by *V. dahliae* strain V991 (LF Zhu, unpublished) against its closely related *V. dahliae* strain CQ2. Subsequently, RNAseq reads overlapping shared core effector genes of *V. dahliae* JR2 and *V. dahliae* strain CQ2 were quantified. We observed that the transcription profiles can be clustered into: 1) transcribed effector genes on both hosts (those with log10 TPM value>0 in both hosts), 2) differentially transcribed effector genes between the two hosts (those with log10 TPM value of 0 in one host and >0 in the other host), and 3) non-transcribed effector genes on either hosts (those with TPM value of 0 in both hosts) (Figure 9A). Of the 165 shared core effector genes between JR2 and CQ2, 61 effector genes were transcribed during cotton as well as *N. benthamiana* colonization, whereas 19 effector genes were only transcribed on cotton, and 41 effector genes were transcribed on *N. benthamiana* (Figure 9B). Additionally, we identified 44 effector genes that were non-transcribed in both strains (Figure 9A). Thus, differential *V. dahliae* core effector gene expression is observed on different host plants. Overall, we are not able to link the composition or the expression of effector gene catalogs to the ability to infect particular host plants.

**Figure 9.**
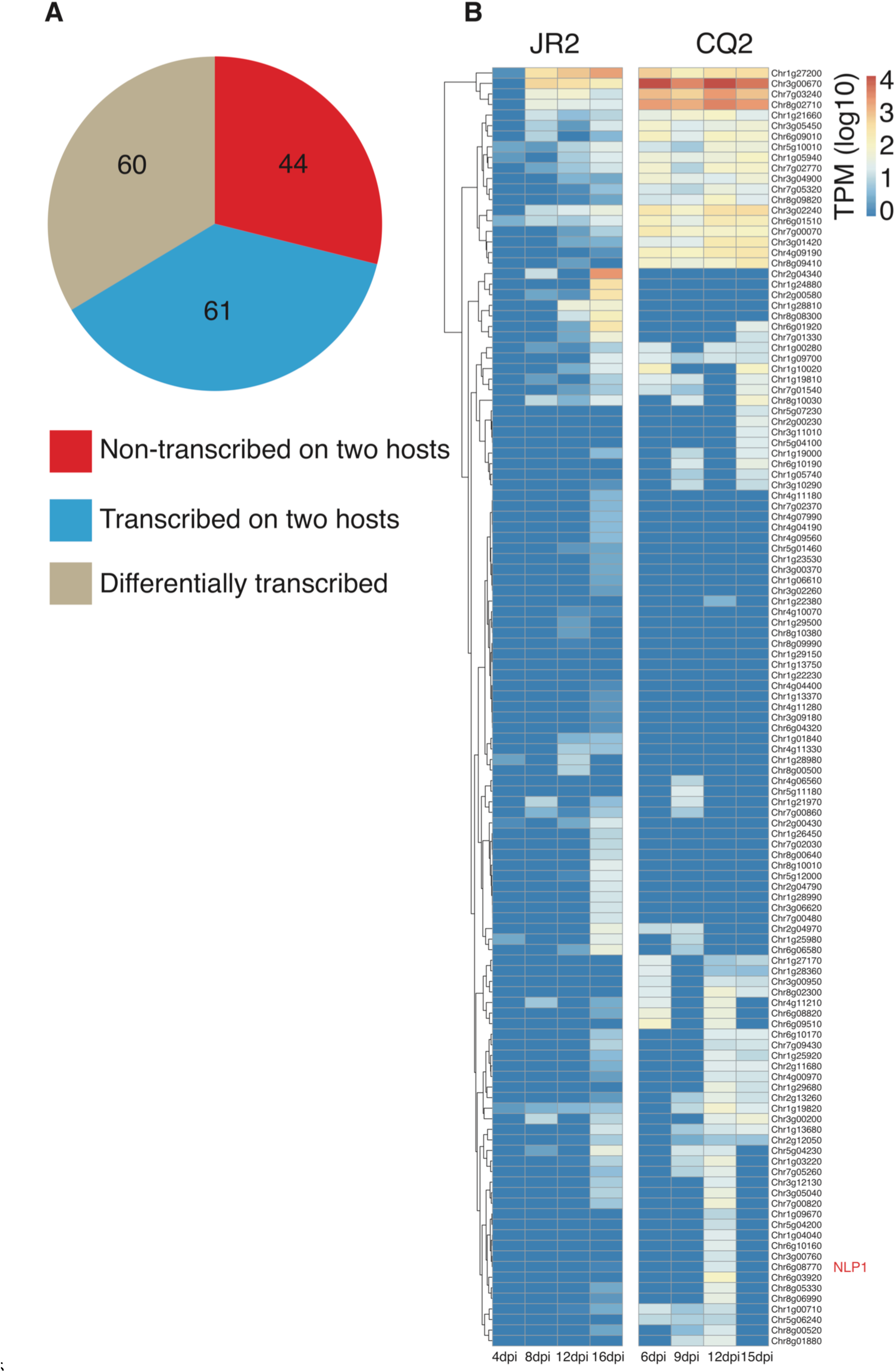
Variability in transcription profiles of core effector genes of *V. dahliae* strains JR2 and CQ2. **A)** Pie chart summarizing the number of non-transcribed, transcribed, and differentially transcribed core effector genes of *V. dahliae* strain JR2 and CQ2 on *N. benthamiana* and cotton, respectively. **B)** Heatmap of the TPM values of core effector genes of *V. dahliae* strain JR2 (left) and *V. dahliae* strain CQ2 (right). The color gradient shows log-scaled TPM values.

## Discussion

For broad host-range pathogens such as *V. dahliae* it is unclear to what extent co-evolution of the pathogen with multiple hosts occurs simultaneously, and what implications this has for their effector repertoires. Whereas it is conceivable that broad host-range pathogens employ core effectors that are active on a multitude of hosts because they target broadly conserved general physiological processes in these hosts, they may also harbor an array of specialized effectors that only exert their activity on particular host species. Presence of such specialized effectors is suggested by the observation that different *V. dahliae* strains, despite their general ability to infect a wide array of host plants, generally display differential capacity to infect particular hosts (Bhat and Subbarao, 1999). Therefore, in this study we investigated whether the absence or presence of particular effectors correlates with the ability to infect particular hosts. First, we predicted the LS effector gene repertoires of *V. dahliae* strains and identified 333 LS effector genes, of which the previously identified LS effector genes *Ave1* of *V. dahliae* strain JR2 (de Jonge et al., 2013) and the *Sun1* effector gene of *V. dahliae* strain 85S (Chapoter 4) were successfully predicted (Figure 8). As improper gene annotation could hamper effector gene discovery (Gibriel et al., 2016), we searched the 333 effector genes against *V. dahliae* genomes using BLAST (tblastn) and successfully identified the previously described effector genes *XLOC_00170, XLOC_008951*, and *XLOC_009059* of *V. dahliae* strain JR2, which are shared by a subset of *V. dahliae* strains (de Jonge et al., 2013) (Figure 8). Intriguingly, we observed that *V. dahliae* strains that are able to infect the same host plant, in this case focused on tomato, cotton, *N. benthamiana*, Arabidopsis, and sunflower, harbor highly divergent LS effector catalogs and not even a single LS effector is shared by the strains that are able to infect either of these host species (Figure 8). This strongly suggests that different strains infect the same host plant by utilizing different effector compositions. If one assumes that different strains of *V. dahliae* must target the same host physiological processes in order to establish the infection, this implies that there must be a significant degree of redundancy among the various effectors that occur in the different strains. Interestingly, fungi of the ascomycete species *Fusarium oxysporum* display a similar infection biology as *V. dahliae*, being soil-borne pathogens that colonize the xylem tissues of their host plants to cause vascular wilt disease and with a largely overlapping host range (Ma et al., 2013). However, whereas individual strains of *V. dahliae* are characterized by their generally broad host range, strains of *F. oxysporum* are generally host-specific and are therefore assigned to *formae speciales* (Ma et al., 2013; Ma et al., 2010). Comparative genomics revealed that *Fusarium* strains able to infect various host plants carry highly overlapping LS effector repertoires (Coleman et al., 2009; Ma et al., 2010; van Dam et al., 2016). Moreover, strains that infect the same host plant cluster based on their LS effector repertoires (Van Dam et al., 2016). Thus, in contrast to *F. oxysporum* strains that harbor highly overlapping LS effectors based on their host range (Coleman et al., 2009; Ma et al., 2010; van Dam et al., 2016), *V. dahliae* strains evolved highly divergent LS effector catalogs, the composition of which does not correlate with the host plant they are able to infect.

Plant pathogens harbor an array of *in planta* highly*-*induced core effector genes that play essential roles on a multitude of hosts (Guyon et al., 2014; Hemetsberger et al., 2015; Santhanam et al., 2013; Yin et al., 2017). We quantified RNAseq reads (RNAseq reads of *V. dahliae* JR2-infecting *N. benthamiana* plants and RNAseq reads of *V. dahliae* strain V991-infecting cotton) overlapping core effector genes of JR2 and CQ2 (a strains closely related to V991). Subsequently, we categorized the effectors in three groups based on their expression profiles. We identified a group of effector genes that are transcribed in both strains (Figure 9B), suggesting that this group of effectors contributes to *V. dahliae* colonization of *N. benthamiana* and cotton. Additionally, we observed a group of effector genes that are highly transcribed in *V. dahliae* strain JR2 on *N. benthamiana* but only lowly transcribed in strain CQ2 on cotton, and vice versa (Figure 9B), suggesting that they differentially contribute to virulence on these two host plants. Furthermore, we identified a group of effector genes that are not transcribed in either strains (Figure 9A), suggesting that this group of core effectors do not play a role in virulence on *N. benthamiana* or on cotton. Nevertheless, it may well be that this group of effector genes is transcribed during *V. dahliae* colonization of other host plants. Surprisingly, we observed that a member of the family of necrosis- and ethylene-inducing-like proteins (NLPs), namely *NLP1*, is not expressed on *N. benthamiana* and is lowly expressed on cotton (Figure 9B), whereas we previously found based on real-time PCR that this effector gene is transcribed in *V. dahliae* strain JR2 on *N. benthamiana* and on tomato, but also in *V. dahliae* strain V592 on cotton (Santhanam et al., 2013; Zhou et al., 2012). Likely, the low expression of *NLP1* expression based on our RNAseq data may be due to the low amount of fungal RNAseq reads among plant-derived ones, as only 0.05% of the reads could be mapped to the *V. dahliae* genome (Faino et al., 2014). Thus, it needs to be taken into account that only highly expressed fungal genes are identified, and it might be worthwhile to confirm whether the group of non-transcribed effector genes still contains lowly or moderately expressed genes based on real-time PCR.

Localization of pathogen effector genes within dynamic genomic regions allows pathogens to rapidly evolve to evade plant immunity once an effector gets recognized by the host (Dong et al., 2015; Raffaele and Kamoun, 2012). When we compared the core and LS effector repertoires between the analysed *V. dahliae* strains, we observed no remarkable differences in the number of core and LS effectors (Table 2). Nevertheless, by comparing features of core and LS effectors we observed that LS effectors are typically shorter in length, localize significantly closer to TEs, and have slightly longer inter-genic lengths when compared with core effectors (Figure 6). Consistent with this observation, a recent study that analysed the genomes of nine different *Verticillium* species showed that species-specific genes displayed significantly shorter gene lengths and longer inter-genic lengths when compared with genes that are conserved across the various species within the genus (Shi-Kunne et al., 2017). Similarly, LS genes of the fungal wheat pathogen *Z. tritici* were frequently found to be shorter, closer to TEs, and have longer inter-genic lengths, when compared with core genes (Haueisen et al., 2017; Plissonneau et al., 2018; Plissonneau et al., 2016). It has been previously suggested that the localization of effector genes in close proximity to TEs and within gene-poor regions mediate rapid evolution of effector catalogs (Raffaele and Kamoun 2012; Seidl and Thomma, 2017). Thus, the localization of LS effector genes of *V. dahliae* strains in close proximity to TEs may mediate accelerated evolution of effector catalogs.

In conclusion, our data demonstrate the extensive variability within the effector repertoires of the broad host range pathogen *V. dahliae*. We have demonstrated that LS effectors are highly divergent among *V. dahliae* strains that infect the same host plant, and core effector genes are differentially expressed between hosts. The variability within LS and core effector genes of *V. dahliae* strains may lead to rapid immunity evasion, which may allow pathogen strains to be competitive in the co-evolution with their multiple hosts.

## Experimental procedures

### *V. dahliae* strains and plant inoculations

In total, 21 *V. dahliae* strains collected at different geographical locations were used in this study (Table S1). All strains were grown on potato dextrose agar (PDA; Oxoid, Basingstoke, UK) at 22°C, and conidiospores were collected from 10-day-old plates and washed with tap water. Disease assays were performed on sunflower (*Helianthus annuus* L. cv. Tutti), cotton (*Gossypium hirsutum* cv. Simian 3), tomato (*Solanum lycopersicum* cv. Moneymaker), *Nicotiana benthamiana* and *Arabidopsis thaliana* (Col-0) plants using the root-dipping inoculation method as previously described (Fradin et al., 2009; Song et al., 2017). Briefly, two-week-old (Arabidopsis, tomato, cotton, sunflower) or three-week-old (*N. benthamiana*) seedlings were carefully uprooted and the roots were rinsed in water. Subsequently, the roots were dipped for eight minutes in a suspension of 10^6^ conidiospores/mL of water. Control plants were treated similarly by root dipping in tap water without conidiospores. Disease symptoms were scored up to 21 (tomato, *N. benthamiana*, Arabidopsis), 28 (cotton) or 45 (sunflower) days post inoculation (dpi).

### Genome sequencing and assembly

The genome sequences of 13 *V. dahliae* strains were previously determined, nine of which were previously sequenced using Illumina HiSeq 2000 (de Jonge et al., 2013; de Jonge et al., 2012) and four that were sequenced using long-read PacBio Single-Molecule Real-Time (SMRT) sequencing technology (Table S1) (Faino et al., 2015). Additionally, eight *V. dahliae* strains (Table S1) were newly sequenced in this study. To this end, genomic DNA of *V. dahliae* strains was obtained from conidiospores that were harvested from ten-day-old cultures grown on potato dextrose agar as described previously (de Jonge et al., 2012). Library preparation (∼500 bp insert size) and genomic sequencing (100 bp paired-end reads) were performed at the Beijing Genome Institute (BGI, Hong Kong). All *V. dahliae* strains that were sequenced with short-read sequencing technology (17 strains) were assembled with the A5 pipeline (default parameters) that automates data cleaning, error correction, assembly, and quality control (Tritt et al., 2012). Genome assembly statistics for all 21 *V. dahliae* strains were calculated using QUAST (Gurevich et al., 2013). Repeats were identified using RepeatModeler (version 1.0.8) (default parameters) (Smit and Hubley, 2010) and masked using RepeatMasker (version 4.0.7) (default parameters) (Smit et al., 2016).

### Gene prediction and annotation

Previously generated gene annotations of *V. dahliae* strains JR2 (Faino et al., 2015) and CQ2 (unpublished) were used in this study. For the remaining 19 *V. dahliae* strains, gene annotation was performed using the Maker2 pipeline (Holt and Yandell, 2011) that combines *ab initio* protein-coding gene evidence from SNAP (Korf 2004), Augustus (Stanke and Waack, 2003), and GeneMark-HMM (Lukashin and Borodovsky, 1998). Additionally, Maker2 was provided with the previously generated reference gene annotation of *V. dahliae* strain JR2 (Faino et al., 2015), gene annotation of *V. dahliae* strain CQ2 (unpublished), and protein homologs of 260 predicted fungal proteomes obtained from the UniProt database (Apweiler et al., 2004).

### Effector profiling

We determined core and LS regions of each *V. dahliae* strain. For LS regions, pairwise whole-genome alignments of the 21 *V. dahliae* strains were performed using NUCmer (version 3.1) (--maxmatch), which is part of the MUMer package (Kurtz et al., 2004), and LS regions (here defined as genomic regions that are shared by <19 *V. dahliae* strains) were extracted. Subsequently, core regions (regions shared by ≥19 *V. dahliae* strains) were determined. Genes localized within core and LS regions were extracted using BEDtools intersect (Quinlan and Hall, 2010).

To identify candidate effectors, N-terminal signal peptides were first predicted with SignalP (version 4.1) (Petersen et al., 2011). Subsequently, the machine-learning approach applied in EffectorP (version 1.0) (default parameters) was used (Sperschneider et al., 2016). Effector genes localized within core and LS regions were extracted using BEDtools intersect (Quinlan and Hall, 2010). Sequence similarity between predicted LS effectors was established by an all-vs.-all analyses using BLASTp (E-value cutoff 1e-5) (Altschul et al., 1990). Clustering of LS effector sequences into different families was performed using MCL (default options) (Li et al., 2003) and visualized using the R package pheatmap (Kolde, 2015).

### Assessment of gene expression

To assess gene expression levels, two RNA-seq datasets were used. The first RNA-seq dataset was previously generated from *V. dahliae* strain JR2-infected *Nicotiana benthamiana* plants at 4, 8, 12, and 16 dpi (de Jonge et al., 2012; Faino et al., 2014). The second RNA-seq dataset was obtained from *V. dahliae* strain V991 infecting *Gossypium hirsutum* (cotton) plants at 6, 9, 12, and 15 dpi (L. Zhu, unpublished). Mapping of RNA-seq datasets to the corresponding genomes was performed using STAR (version 2.5.3) (--runThreadN 16) (Dobin et al., 2013), and gene expression levels were determined using RSEM (version 1.2.3) (calculate-expression command) (default parameters) (Li and Dewey 2011), by calculating transcripts per million (TPM) for each gene in each sample.

## Acknowledgements

J Li acknowledges a PhD fellowship from China Scholarship Council (CSC). Work in the laboratories of M.F.S and B.P.H.J.T. and is supported by the Research Council Earth and Life Sciences (ALW) of the Netherlands Organization of Scientific Research (NWO). The authors declare no conflict of interest.

## References

Altschul, S. F., Gish, W., Miller, W., Myers, E. W., and Lipman, D. J. (1990) Basic local alignment search tool. Journal of Molecular Biology 215.

Apweiler, R., Bairoch, A., Wu, C. H., Barker, W. C., Boeckmann, B., Ferro, S., Gasteiger, E., Huang, H., Lopez, R., Magrane, M., Martin, M. J., Natale, D. A., O’Donovan, C., Redaschi, N., and Yeh, L.-S. L. (2004) Uniprot: the universal protein knowledgebase. Nucleic Acids Research 32:D115–D119.

Bhat, R. G., and Subbarao, K. V. (1999) Host range specificity in Verticillium dahliae. Phytopathology 89:1218–1225.

Coleman, J. J., Rounsley, S. D., Rodriguez-Carres, M., Kuo, A., Wasmann, C. C., Grimwood, J., Schmutz, J., Taga, M., White, G. J., Zhou, S., Schwartz, D. C., Freitag, M., Ma, L.-j., Danchin, E. G. J., Henrissat, B., Coutinho, P. M., Nelson, D. R., Straney, D., Napoli, C. A., Barker, B. M., Gribskov, M., Rep, M., Kroken, S., Molnár, I., Rensing, C., Kennell, J. C., Zamora, J., Farman, M. L., Selker, E. U., Salamov, A., Shapiro, H., Pangilinan, J., Lindquist, E., Lamers, C., Grigoriev, I. V., Geiser, D. M., Covert, S. F., Temporini, E., and VanEtten, H. D. (2009) The genome of Nectria haematococca: Contribution of supernumerary chromosomes to gene expansion. PLoS Genetics 5:e1000618.

Cook, D. E., Mesarich, C. H., and Thomma, B. P. H. J. (2015) Understanding plant immunity as a surveillance system to detect invasion. Annual Review of Phytopathology 53:541–563.

Croll, D., and McDonald, B. A. (2012) The accessory genome as a cradle for adaptive evolution in pathogens. PLoS Pathogens 8:e1002608.

de Jonge, R., Bolton, M. D., Kombrink, A., van den Berg, G. C. M., Yadeta, K. A., and Thomma, B. P. H. J. (2013) Extensive chromosomal reshuffling drives evolution of virulence in an asexual pathogen. Genome Research 23:1271–1282.

de Jonge, R., Bolton, M. D., and Thomma, B. P. H. J. (2011) How filamentous pathogens co-opt plants: the ins and outs of fungal effectors. Current Opinion in Plant Biology 14:400–406.

de Jonge, R., Peter van Esse, H., Maruthachalam, K., Bolton, M. D., Santhanam, P., Saber, M. K., Zhang, Z., Usami, T., Lievens, B., Subbarao, K. V., and Thomma, B. P. H. J. (2012) Tomato immune receptor Ve1 recognizes effector of multiple fungal pathogens uncovered by genome and RNA sequencing. Proceedings of the National Academy of Sciences of the United States of America 109:5110–5115.

de Jonge, R., van Esse, H. P., Kombrink, A., Shinya, T., Desaki, Y., Bours, R., van der Krol, S., Shibuya, N., Joosten, M. H., and Thomma, B. P. H. J. (2010) Conserved fungal LysM effector Ecp6 prevents chitin-triggered immunity in plants. Science 329:953–955.

Dobin, A., Davis, C. A., Schlesinger, F., Drenkow, J., Zaleski, C., Jha, S., Batut, P., Chaisson, M., and Gingeras, T. R. (2013) STAR: ultrafast universal RNA-seq aligner. Bioinformatics 29:15–21.

Dong, S., Raffaele, S., and Kamoun, S. (2015) The two-speed genomes of filamentous pathogens: waltz with plants. Current Opinion in Genetics & Development 35:57–65.

Faino, L., de Jonge, R., and Thomma, B. P. H. J. (2014) The transcriptome of Verticillium dahliae-infected Nicotiana benthamiana determined by deep RNA sequencing. Plant Signaling & Behavior 7:1065–1069.

Faino L, Seidl MF, Datema E, van den Berg GC, Janssen A, Wittenberg AH, Thomma BPHJ (2015) Single-molecule real-time sequencing combined with optical mapping yields completely finished fungal genome. mBio 6: e00936–00915.

Fisher, M. C., Henk, D. A., Briggs, C. J., Brownstein, J. S., Madoff, L. C., McCraw, S. L., and Gurr, S. J. (2012) Emerging fungal threats to animal, plant and ecosystem health. Nature 484:186–194.

Fradin, E. F., and Thomma, B. P. H. J. (2006) Physiology and molecular aspects of Verticillium wilt diseases caused by V. dahliae and V. alboatrum. Molecular Plant Pathology 7:71–86.

Fradin, E. F., Zhang, Z., Juarez Ayala, J. C., Castroverde, C. D. M., Nazar, R. N., Robb, J., Liu, C.-M., and Thomma, B. P. H. J. (2009) Genetic dissection of Verticillium wilt resistance mediated by tomato Ve1. Plant Physiology 150:320–332.

Gibriel, H. A. Y., Thomma, B. P. H. J., and Seidl, M. F. (2016) The age of effectors: genome-based discovery and applications. Phytopathology 106: 1206–1212.

Gurevich, A., Saveliev, V., Vyahhi, N., and Tesler, G. (2013) QUAST: quality assessment tool for genome assemblies. Bioinformatics 29:1072– 1075.

Guyon, K., Balagué, C., Roby, D., and Raffaele, S. (2014) Secretome analysis reveals effector candidates associated with broad host range necrotrophy in the fungal plant pathogen Sclerotinia sclerotiorum. BMC Genomics 15:336.

Haueisen, J., Moeller, M., Eschenbrenner, C. J., Grandaubert, J., Seybold, H., Adamiak, H., and Stukenbrock, E. H. (2017) Extremely flexible infection programs in a fungal plant pathogen. bioRxiv: https://doi.org/10.1101/229997.

Hemetsberger, C., Mueller, A. N., Matei, A., Herrberger, C., Hensel, G., Kumlehn, J., Mishra, B., Sharma, R., Thines, M., Hückelhoven, R., and Doehlemann, G. (2015) The fungal core effector Pep1 is conserved across smuts of dicots and monocots. New Phytologist 206:1116–1126.

Holt, C., and Yandell, M. (2011) MAKER2: an annotation pipeline and genome-database management tool for second-generation genome projects. BMC Bioinformatics 12:491.

Inderbitzin, P., Bostock, R. M., Davis, R. M., Usami, T., Platt, H. W., and Subbarao, K. V. (2011) Phylogenetics and taxonomy of the fungal vascular wilt pathogen Verticillium, with the descriptions of five new species. PLoS One 6:e28341.

Jones, J. D. G., and Dangl, J. L. (2006) The plant immune system. Nature 444:323–329.

Kema, G. H. J., Mirzadi Gohari, A., Aouini, L., Gibriel, H. A. Y., Ware, S. B., van den Bosch, F., Manning-Smith, R., Alonso-Chavez, V., Helps, J., Ben M’Barek, S., Mehrabi, R., Diaz-Trujillo, C., Zamani, E., Schouten, H. J., van der Lee, T. A. J., Waalwijk, C., de Waard, M. A., De Wit, P. J. G. M., Verstappen, E. C. P., Thomma, B. P. H. J., Meijer, H. J. G., and Seidl, M. F. (2018) Stress and sexual reproduction affect the dynamics of the wheat pathogen effector AvrStb6 and strobilurin resistance. Nature Genetics 23:678.

Kolde, R. 2015. pheatmap: Pretty Heatmaps. R package version 1.0. 8. Kombrink, A., Rovenich, H., Shi-Kunne, X., Rojas-Padilla, E., van den Berg, G. C. M., Domazakis, E., de Jonge, R., Valkenburg, D.-J., Sánchez-Vallet, A., Seidl, M. F., and Thomma, B. P. H. J. (2017) Verticillium dahliae LysM effectors differentially contribute to virulence on plant hosts. Molecular Plant Pathology 18:596–608.

Korf, I. (2004) Gene finding in novel genomes. BMC Bioinformatics 5: 59.

Kurtz, S., Phillippy, A., Delcher, A. L., Smoot, M., Shumway, M., Antonescu, C., and Salzberg, S. L. (2004) Versatile and open software for comparing large genomes. Genome Biology 5: R12.

Li, B., and Dewey, C. N. (2011) RSEM: accurate transcript quantification from RNA-Seq data with or without a reference genome. BMC Bioinformatics 12:323.

Li, L., Stoeckert, C. J., and Roos, D. S. (2003) OrthoMCL: identification of ortholog groups for eukaryotic genomes. Genome Research 13:2178– 2189.

Lukashin, A. V., and Borodovsky, M. (1998) GeneMark.hmm: new solutions for gene finding. Nucleic Acids Research 26.

Ma, L.-J., Geiser, D. M., Proctor, R. H., Rooney, A. P., O’Donnell, K., Trail, F., Gardiner, D. M., Manners, J. M., and Kazan, K. (2013) Fusarium pathogenomics. Annual Review of Microbiology 67:399–416.

Ma, L. J., Does, H. C., Borkovich, K. A., Coleman, J. J., Daboussi, M. J., Pietro, A., Dufresne, M., Freitag, M., Grabherr, M., and Henrissat, B. (2010) Comparative genomics reveals mobile pathogenicity chromosomes in Fusarium. Nature 464.

Marshall, R., Kombrink, A., Motteram, J., Loza-Reyes, E., Lucas, J., Hammond-Kosack, K. E., Thomma, B. P. H. J., and Rudd, J. J. (2011) Analysis of two in planta expressed LysM effector homologs from the fungus Mycosphaerella graminicola reveals novel functional properties and varying contributions to virulence on wheat. Plant Physiology 156:756–769.

Meile, L., Croll, D., Brunner, P. C., Plissonneau, C., Hartmann, F. E., McDonald, B. A., and Sánchez-Vallet, A. (2018) A fungal avirulence factor encoded in a highly plastic genomic region triggers partial resistance to septoria tritici blotch. New Phytologist 65:512.

Mentlak, T. A., Kombrink, A., Shinya, T., Ryder, L. S., Otomo, I., Saitoh, H., Terauchi, R., Nishizawa, Y., Shibuya, N., Thomma, B. P. H. J., and Talbot, N. J. (2012) Effector-mediated suppression of chitin-triggered immunity by Magnaporthe oryzae is necessary for rice blast disease. The Plant Cell 24:322–335.

Pennisi, E. (2010) Armed and dangerous. Science 327:804–805.

Petersen, T. N., Brunak, S., von Heijne, G., and Nielsen, H. (2011) SignalP 4.0: discriminating signal peptides from transmembrane regions. Nature Methods 8:785–786.

Plissonneau, C., Hartmann, F. E., and Croll, D. (2018) Pangenome analyses of the wheat pathogen Zymoseptoria tritici reveal the structural basis of a highly plastic eukaryotic genome. BMC Biology 16:673.

Plissonneau, C., Stürchler, A., and Croll, D. (2016) The evolution of orphan regions in genomes of a fungal pathogen of wheat. mBio 7.

Quinlan, A. R., and Hall, I. M. (2010) BEDTools: a flexible suite of utilities for comparing genomic features. Bioinformatics 26.

Raffaele, S., and Kamoun, S. (2012) Genome evolution in filamentous plant pathogens: why bigger can be better. Nature Review Microbiology 10:417–430.

Santhanam, P., van Esse, H. P., Albert, I., Faino, L., Nürnberger, T., and Thomma, B. P. H. J. (2013) Evidence for functional diversification within a fungal NEP1-like protein family. Molecular Plant Microbe Interactions 26:278–286.

Schmidt, S. M., Houterman, P. M., Schreiver, I., Ma, L., Amyotte, S., Chellappan, B., Boeren, S., Takken, F. L. W., and Rep, M. (2013) MITEs in the promoters of effector genes allow prediction of novel virulence genes in Fusarium oxysporum. BMC Genomics 14:1–21.

Seidl, M. F., and Thomma, B. (2017a.) Transposable elements direct the coevolution between plants and microbes. Trends in Genetics 33:842–851.

Shi-Kunne, X., Faino, L., van den Berg, G. C. M., Thomma, B. P. H. J., and Seidl, M. F. (2017) Evolution within the fungal genus Verticillium is characterized by chromosomal rearrangement and gene loss. Environmental Microbiology 20:1362–1373.

Simão, F. A., Waterhouse, R. M., Ioannidis, P., Kriventseva, E. V., and Zdobnov, E. M. (2015) BUSCO: assessing genome assembly and annotation completeness with single-copy orthologs. Bioinformatics 31:3210–3212.

Smit, A., and Hubley, R. (2010) RepeatModeler Open-1.0. Repeat Masker Website. http://www.repeatmasker.org/Repeat-Modeler.html.

Smit, A., Hubley, R., and Green, P. (2016) RepeatMasker Open-4.0. Available from: http://www.repeatmasker.org

Song, Y., Liu, L., Wang, Y., Valkenburg, D.-J., Zhang, X., Zhu, L., and Thomma, B. P. H. J. (2017) Transfer of tomato immune receptor Ve1 confers Ave1-dependent Verticillium resistance in tobacco and cotton. Plant Biotechnology Journal 16:638–648.

Sperschneider, J., Gardiner, D. M., Dodds, P. N., Tini, F., Covarelli, L., Singh, K. B., Manners, J. M., and Taylor, J. M. (2016) EffectorP: predicting fungal effector proteins from secretomes using machine learning. New Phytologist 210:743–761.

Stanke, M., and Waack, S. (2003) Gene prediction with a hidden Markov model and a new intron submodel. Bioinformatics 19.

Stergiopoulos, I., van den Burg, H. A., Okmen, B., Beenen, H. G., van Liere, S., Kema, G. H., and de Wit, P. J. (2010) Tomato Cf resistance proteins mediate recognition of cognate homologous effectors from fungi pathogenic on dicots and monocots. Proceedings of the National Academy of Sciences of the United States of America 107:7610– 7615.

Takahara, H., Hacquard, S., Kombrink, A., Hughes, H. B., Halder, V., Robin, G. P., Hiruma, K., Neumann, U., Shinya, T., Kombrink, E., Shibuya, N., Thomma, B. P. H. J., and O’Connell, R. J. (2016) Colletotrichum higginsianum extracellular LysM proteins play dual roles in appressorial function and suppression of chitin-triggered plant immunity. New Phytologist 211:1323–1337.

Tritt, A., Eisen, J. A., Facciotti, M. T., and Darling, A. E. (2012) An integrated pipeline for de novo assembly of microbial genomes. PLoS One 7:e42304.

van Dam, P., Fokkens, L., Schmidt, S. M., Linmans, J. H. J., Kistler, H. C., Ma, L.-J., and Rep, M. (2016) Effector profiles distinguish formae speciales of Fusarium oxysporum. Environmental Microbiology 18:4087–4102.

Yin, J., Gu, B., Huang, G., Tian, Y., Quan, J., Lindqvist-Kreuze, H., and Shan, W. (2017) Conserved RXLR effector genes of Phytophthora infestans expressed at the early stage of potato infection are suppressive to host defense. Frontiers in Plant Sciences 8.

Zhou, B. J., Jia, P. S., Gao, F., and Guo, H. S. (2012) Molecular characterization and functional analysis of a necrosis- and ethylene-inducing, protein-encoding gene family from Verticillium dahliae. Molecular Plant-Microbe Interactions 25:964–975.

