## Supplemental Data for "*Verticillium dahliae* strains that infect the same host plant display highly divergent effector catalogs"

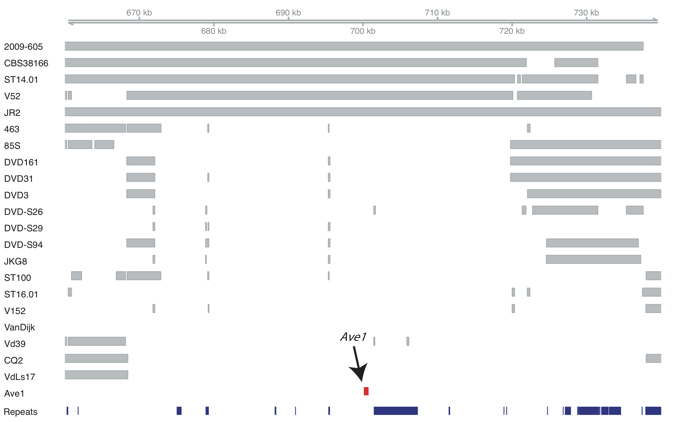

**Figure S1. *V. dahliae* effector gene *Ave1* is localized within a lineage-specific region.** A highly variable, repeat-rich (blue), lineage-specific (LS) region, which harbors the *Ave1* effector gene (red) of *V. dahliae* strain JR2 is shown. This LS effector gene is only present in a subset of *V. dahliae* strains. Grey bars indicate the genome alignments of *V. dahliae* strains to the reference strain JR2. This effector gene is only present in strains 2009-605, CBS38166, ST14.01, V52, and JR2, but absent in the other strains.

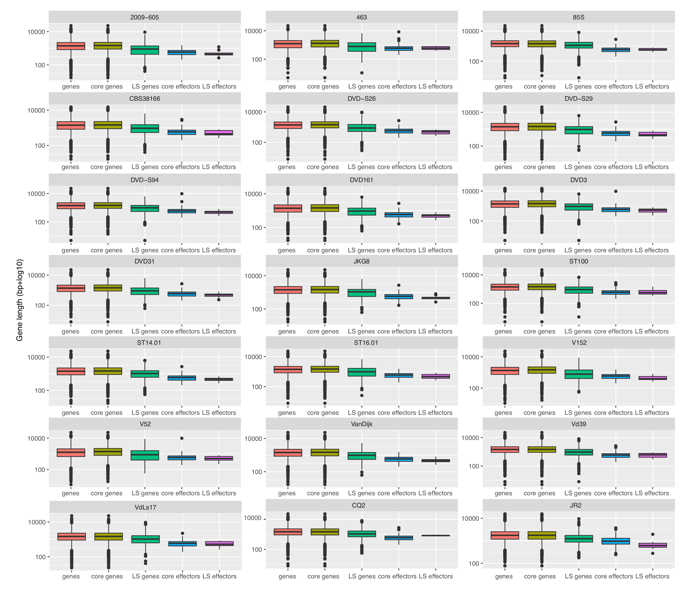

**Figure S2.** **Length of genes and effector genes of *V. dahliae* strains.** Gene length (log10) is shown for all genes, core genes, LS genes, core effectors, and LS effectors for all *V. dahliae* strains.

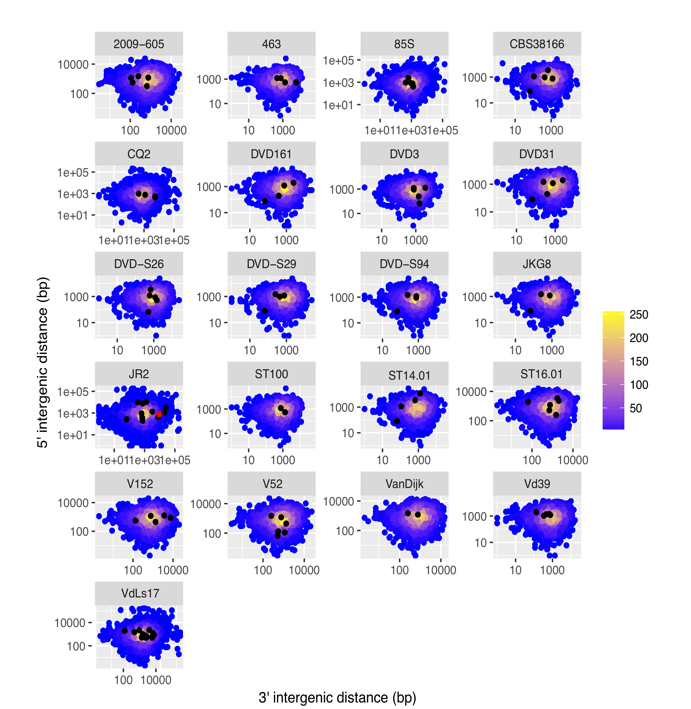

**Figure S3.** **5′ and 3′ inter-genic length of genes and effector genes of *V. dahliae* strains.** Inter-genic length (bp) is shown for all genes (blue) and LS effector genes (black) for all *V. dahliae* strains. *Ave1* effector gene of *V. dahliae* strain JR2 is highlighted in red.

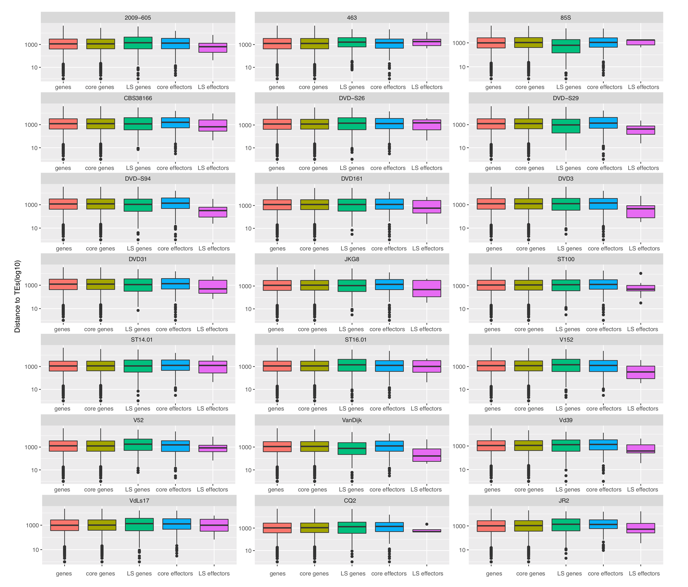

**Figure S4.** **Distance to TEs of genes and effector gnenes of *V. dahliae* strains.** Distance to TEs (log10) is shown for all genes, core genes, LS genes, core effectors, and LS effectors for all *V. dahliae* strains.

**Table S1. *Verticillium dahliae* strains used in this study.**

| Strain | Sequencing platform | Reference | Originating host | Geographical location |
| --- | --- | --- | --- | --- |
| CQ2 | PacBio | (Depotter et al., 2018) | Cotton | China |
| 85S | PacBio | (Depotter et al., 2018) | Sunflower | France |
| VdLs17 | PacBio | (Faino et al., 2015) | Lettuce | Ca, USA |
| JR2 | PacBio | (Faino et al., 2015) | Tomato | ON, Canada |
| CBS38166 | Illumina HiSeq 2000 | (de Jonge et al., 2012) | Tomato | QC, Canada |
| ST14.01 | Illumina HiSeq 2000 | (de Jonge et al., 2012) | Pistachio | CA, USA |
| ST100 | Illumina HiSeq 2000 | (de Jonge et al., 2013) | Soil | Belgium |
| DVD3 | Illumina HiSeq 2000 | (de Jonge et al., 2012) | Potato | Canada |
| DVD31 | Illumina HiSeq 2000 | (de Jonge et al., 2012) | Tomato | Canada |
| DVD161 | Illumina HiSeq 2000 | (de Jonge et al., 2012) | Potato | ON, Canada |
| DVD-S26 | Illumina HiSeq 2000 | (de Jonge et al., 2012) | Soil | Canada |
| DVD-S29 | Illumina HiSeq 2000 | (de Jonge et al., 2012) | Soil | Canada |
| DVD-S94 | Illumina HiSeq 2000 | (de Jonge et al., 2012) | Soil | Canada |
| JKG8 | Illumina HiSeq 2000 | This study | Potato | The Netherlands |
| 2009-605 | Illumina HiSeq 2000 | This study | Bell pepper | Ukraine |
| 463 | Illumina HiSeq 2000 | This study | Cotton | Mexico |
| ST16.01 | Illumina HiSeq 2000 | This study | Cotton | Syria |
| V152 | Illumina HiSeq 2000 | This study | Oak | Hungry |
| V52 | Illumina HiSeq 2000 | This study | Pepper | Austria |
| Vd39 | Illumina HiSeq 2000 | This study | Sunflower | Germany |
| VanDijk | Illumina HiSeq 2000 | This study | Chrysanthemum | The Netherlands |

**Table S2**. **Assembly statistics for the various *Verticillium* *dahliae* genomes.**

| Strain | Genome size (Mb) | # scaffolds (>= 0 bp) | # scaffolds (>= 1000 bp) | GC (%) | N50 (Kb) | # N's per 100 kbp | BUSCO (%) |
| --- | --- | --- | --- | --- | --- | --- | --- |
| 2009-605 | 34.06 | 1931 | 1521 | 54.76 | 55.11 | 123.45 | 95.4 |
| 463 | 34.03 | 4188 | 3562 | 53.47 | 17.74 | 244.06 | 75 |
| 85S | 35.93 | 40 | 40 | 53.55 | 3176.09 | 0 | 99.3 |
| CBS38166 | 34.03 | 2092 | 1727 | 54.24 | 45.08 | 316.06 | 91.6 |
| DVD161 | 33.47 | 2155 | 1819 | 54.4 | 41.42 | 330.79 | 90.8 |
| DVD31 | 33.58 | 2429 | 2064 | 54.14 | 35.75 | 336.71 | 89.2 |
| DVD3 | 34.42 | 1921 | 1693 | 53.62 | 42.77 | 378.65 | 88.9 |
| DVD-S26 | 34.74 | 2275 | 1894 | 54.2 | 43.54 | 367.67 | 92.2 |
| DVD-S29 | 33.14 | 2226 | 1811 | 54.56 | 42.73 | 306.89 | 91.6 |
| DVD-S94 | 34.42 | 1730 | 1494 | 53.92 | 53.17 | 287 | 91.9 |
| JKG8 | 33.85 | 1840 | 1458 | 54.5 | 56.35 | 172.12 | 95.6 |
| ST100 | 34.92 | 2137 | 1756 | 53.63 | 49.89 | 315.39 | 93 |
| ST14.01 | 34.48 | 1571 | 1336 | 54.03 | 61.81 | 221.84 | 95 |
| ST16.01 | 34.22 | 1821 | 1454 | 54.9 | 57.46 | 121.71 | 95.6 |
| V152 | 33.98 | 2539 | 2174 | 54.28 | 32.47 | 146.98 | 87.6 |
| V52 | 33.54 | 3419 | 2920 | 54.39 | 21.80 | 162.37 | 78.6 |
| VanDijk | 33.17 | 1769 | 1389 | 54.71 | 61.05 | 109.26 | 95.7 |
| Vd39 | 35.90 | 1579 | 1222 | 53.55 | 94.44 | 222.52 | 98.7 |
| CQ2 | 35.82 | 17 | 17 | 53.26 | 3754.19 | 0 | 97.5 |
| JR2 | 36.15 | 8 | 8 | 53.89 | 4168.63 | 0 | 99.4 |
| VdLs17 | 35.97 | 8 | 8 | 54 | 5894.01 | 0 | 98.9 |

**Table S3. Summary of transposable elements of the various *Verticillium dahliae* strains.**

|  | SINEs | | LINEs | | LTRs | |
| --- | --- | --- | --- | --- | --- | --- |
| Strain | length occupied (bb) | percentage of sequence (%) | length occupied (bb) | percentage of sequence (%) | length occupied (bb) | percentage of sequence (%) |
| 2009-605 | 0 | 0 | 0 | 0 | 1403090 | 4.12 |
| 463 | 0 | 0 | 43660 | 0.13 | 1601065 | 4.7 |
| 85S | 2289.00 | 0.01 | 57668.00 | 0.16 | 3082210.00 | 8.58 |
| CBS38166 | 0.00 | 0.00 | 29731.00 | 0.09 | 1625629.00 | 4.78 |
| DVD161 | 1994.00 | 0.01 | 0.00 | 0.00 | 1871766.00 | 5.59 |
| DVD31 | 15326.00 | 0.05 | 14424.00 | 0.04 | 1718276.00 | 5.12 |
| DVD3 | 1923.00 | 0.01 | 17323.00 | 0.05 | 2262138.00 | 6.57 |
| DVD-S26 | 0.00 | 0.00 | 34185.00 | 0.10 | 1847224.00 | 5.32 |
| DVD-S29 | 22393.00 | 0.01 | 7832.00 | 0.02 | 1297199.00 | 3.91 |
| DVD-S94 | 0.00 | 0.00 | 23539.00 | 0.07 | 2082451.00 | 6.05 |
| JKG8 | 2294.00 | 0.01 | 0.00 | 0.00 | 1344573.00 | 3.97 |
| ST100 | 1720.00 | 0.01 | 50597.00 | 0.14 | 2693973.00 | 7.71 |
| ST14.01 | 0.00 | 0.00 | 30264.00 | 0.09 | 1610719.00 | 4.67 |
| ST16.01 | 1582.00 | 0.00 | 50167.00 | 0.15 | 1009974.00 | 2.95 |
| V152 | 0.00 | 0.00 | 40614.00 | 0.12 | 1660694.00 | 4.89 |
| V52 | 3965.00 | 0.01 | 28553.00 | 0.09 | 1442720.00 | 4.30 |
| VanDijk | 1954.00 | 0.01 | 0 | 0 | 1337583.00 | 4.03 |
| Vd39 | 1582.00 | 0.00 | 71392.00 | 0.20 | 3201873.00 | 8.92 |
| CQ2 | 3965.00 | 0.01 | 54386.00 | 0.15 | 3156817.00 | 8.81 |
| JR2 | 1954.00 | 0.01 | 98739.00 | 0.27 | 2841008.00 | 7.86 |
| VdLs17 | 0.00 | 0.00 | 235025.00 | 0.65 | 2875975.00 | 7.99 |
